# Phantom limb pain intensity is associated with generalized hyperalgesia

**DOI:** 10.1101/538207

**Authors:** Xaver Fuchs, Martin Diers, Jörg Trojan, Pinar Kirsch, Christopher Milde, Robin Bekrater-Bodmann, Mariela Rance, Jens Foell, Jamila Andoh, Susanne Becker, Herta Flor

**Affiliations:** Biopsychology and Cognitive Neuroscience, Faculty of Psychology and Sports Science, Bielefeld University, Bielefeld, Germany; Department of Cognitive and Clinical Neuroscience, Central Institute of Mental Health, Medical Faculty Mannheim, Heidelberg University, Mannheim, Germany; Department of Psychosomatic Medicine and Psychotherapy, LWL University Hospital, Ruhr-University Bochum, Bochum, Germany; Department of Psychology, University of Koblenz-Landau, Landau, Germany; Department of Radiology and Biomedical Imaging, Yale School of Medicine, New Haven, CT, USA; Department of Psychology, Florida State University, Tallahassee, Florida, USA

## Abstract

After limb amputation, most amputees suffer from phantom limb pain (PLP). The mechanisms underlying this condition are complex and insufficiently understood. Altered somatosensory sensitivity (either heightened or lowered) might contribute to PLP. Recent studies have tested this assumption but mainly focused on the residual limb. However, altered somatosensation in PLP might be generalized. In this study, we applied quantitative sensory testing to 37 unilateral upper-limb amputees (23 with PLP, 14 without PLP) and 19 healthy controls. We assessed thresholds to heat pain (HPT), pressure pain (PPT), warmth detection (WDT), and two-point discrimination (2PDT) at the residual limb, a homologous point and the thenar of the intact limb, and both corners of the mouth. We did not find significant differences in any of the thresholds between the groups. However, higher PLP intensity was significantly related to lower HPT at all measured body sites except for the residual limb. At the residual limb, lower HPT were observed in more distal amputations and in amputees showing a higher degree of prosthesis use. Although WDT did by itself not significantly correlate with PLP intensity at any of the body sites, multiple regression analysis showed the highest multiple correlations with PLP intensity for a combination of high WDT and low HPT at the corners of the mouth. In this model, the combination of HPT and WDT shared 58% of the variance with PLP intensity. Other factors of potential importance, especially residual limb pain, were not significantly associated to any sensory threshold. We conclude that the intensity, but not the presence of PLP is positively associated with higher heat pain sensitivity. Since this association was observed at various, distributed body sites, we suggest that central mechanisms might be underlying such generalized hyperalgesia.

## 1. Introduction

Phantom limb pain (PLP)—the sensation of pain located outside one’s physical body in a lost limb—is a common consequence of amputation and occurs in about 50–80%^52^ of all amputees. Both peripheral and central mechanisms are involved in chronic PLP^18,58^. However, their exact roles and impact are under debate. Peripheral factors relate to nociceptive input stemming from the residual limb and can involve peripheral sensitization, spontaneous discharges from the residual limb due to involuntary movement or physiological arousal^60^, neuromas^10,21,67^, or activity generated in dorsal root ganglia^64^. Central sensitization can develop as a consequence of long-term peripheral nociceptive input and is mediated by spinal cord neurons^38,72^. In the brain, PLP has been associated with maladaptive reorganization of body maps in the primary somatosensory (SI)^16^ and primary motor cortex^42^ as well as preserved function of the phantom cortex^49,35^.

Increased pain sensitivity encompasses both peripheral and central mechanisms^18^ and is commonly assessed psychophysically by quantitative sensory testing (QST)^46,47,59^. Although QST has previously been applied in amputees, the results are mixed and do not clearly support an association with PLP. This might be related to methodological differences between studies. Most previous studies compared the residual limb to a corresponding area on the intact limb. Some previous studies reported enhanced tactile sensitivity^27,31,61,68^, other studies reported higher sensitivity of the residual limb to thermal^14,65^ or pressure pain^7,36,63^, but did not report whether these alterations in sensitivity were related to PLP. Other studies examined the relationship of tactile and pain sensitivity to PLP, but most did not find a significant relationship with tactile sensitivity^32,25,30,41^ or with thermal pain^17,28,30,41^. One study reported hypoesthesia to cold stimuli at the residual limb, which tended to be significantly correlated with PLP intensity^28^. Another study found hyperalgesia for electrical pain stimuli at the intact limb in amputees with but not in those without PLP, suggesting central sensitization in PLP^41^.

Inconsistent findings of previous studies might partly be explained by small sample sizes and heterogeneity of the samples (e.g. lower-limb vs. upper-limb amputations) and methods. In particular, comparisons relating the residual to the intact limb might be problematic since it is difficult to standardize measures taken at residual limbs due to variations in levels of amputation and scar tissue. QST of the residual limb may relate to residual limb pain (RLP) rather than PLP. RLP might also be associated with peripheral sensitization, although empirical evidence is lacking. Testing only the residual limb and a homologous contralateral site cannot determine whether pain sensitivity is also present at remote sites, for instance the mouth in upper limb amputees, which is adjacent to the cortical representation of the phantom limb and might have altered sensitivity related to reorganization or due to central sensitization^70–72^.

In order to clarify the possibility that changes might be more generalized, the present study tested whether PLP is accompanied by changes in sensitivity to noxious and non-noxious stimuli applied at several standardized body sites. The aims were (a) to investigate whether there are changes in somatosensory thresholds, which are associated with the presence and/or the intensity of PLP, and (b) to test whether potential changes are restricted to the residual limb, which could involve peripheral mechanisms, and/or whether they are present in other areas of the body, which could point at central mechanisms. Finally, we sought to (c) take into account asscociations with other variables which are potential drivers of sensitization and brain changes, especially RLP and prosthesis use^33,43^.

## 2. Methods

### 2.1. Participants

Thirty-seven unilateral upper-limb amputees and 19 healthy controls (HC) took part in this study. We employed QST and the assessment of phantom phenomena and chronic pain. The sample was recruited from a nation-wide data base we had previously collected^4^. The study was carried out at the Central Institute of Mental Health in Mannheim, Germany, and was approved by the Medical Ethics Commission II of the Medical Faculty Mannheim, Heidelberg University. Amputees were divided into two groups—one with PLP (PLP; N=23) and one without PLP (nonPLP; N=14). Grouping was based on the “phantom pain severity” subscale of the German version of the West Haven-Yale Multidimensional Pain Inventory^16,19,34^ (MPI) adapted for separate assessments of phantom and residual limb pain. The MPI is a 22-item questionnaire that uses a 7-point numeric rating scale for each item ranging from 0 to 6. The “pain severity” subscale is calculated as the average value from three items assessing (1) the intensity of momentary pain, (2) the average pain during the last week and (3) the intensity of suffering from pain. Amputees with a score of 0 related to PLP were assigned to the nonPLP, all other amputees to the PLP group. The HC were matched for age, sex and stimulation sites (see below) with amputees from both the PLP or the nonPLP group. A description of the participants and of the matching between amputees and HC is provided in Table 1.

There were no statistically significant differences between the three groups (PLP, nonPLP and HC) in the distribution of sex (χ^2^_(2, *N*=56)_ = 0.06, *p* = 0.97) and age (*F*_(2, 53)_ = 0.10, *p* = 0.90). The PLP group and the nonPLP group did not significantly differ in their level of amputation (*t*_(22.7)_ = −1.1, *p* = 0.28) or side of amputation (χ^2^_(1, *N*=37)_ = 0.53, *p* = 0.47). The PLP group was amputated at a higher age than the nonPLP group (27.1 vs. 19.8 years), which tended to be significant (*t*_(29.5)_ = 1.8, *p* = 0.08). All amputees were right-handed before amputation and so were all of the HC, according to the Edinburgh Handedness Inventory^55^.

**Table 1:** Description of the study sample. Used abbreviations: PLP = phantom limb pain, nonPLP = without phantom limb pain, HC = healthy control, RLP = residual limb pain, EHI = Edinburgh handedness inventory, myo-el. = myoelectric prosthesis. ^1^A value of 100 in the EHI indicates the strongest degree of right-handedness, a value of −100 strongest degree of left-handedness. ^2^Level of stimulation was defined as the position of measurements at the arm relative to the length of the (intact) arm, measured from the caput humeri to the tip of the third digit. In the amputees, this position was defined by the length of the residual limb. Homologous sites were measured in the matched HC. ^3^Side of amputation in the HC (in brackets) refers to the side of amputation of the matched amputee. ^4^The category “other” causes of amputation subsumes all causes not falling into the categories traumatic, injury, dysplasia, infection, tumor or vascular disease. ^5^Pain severity subscale of the MPI adapted for PLP and RLP respectively. Values range from 0 (lowest severity) to 6 (highest severity).

| ID | Group | Mat-<br>ching | Sex | Age | EHI<br>score<sup>1</sup> | Lvl. of<br>stim.<br>(%)<sup>2</sup> | Age at<br>amp. | Side of<br>amp.<sup>3</sup> | Cause of<br>amp.<sup>4</sup> | Prosthesis | Phan-<br>tom<br>limb | Tele-<br>scope | PLP<br>intensity<sup>5</sup> | RLP<br>intensity<sup>5</sup> |
| --- | --- | --- | --- | --- | --- | --- | --- | --- | --- | --- | --- | --- | --- | --- |
| A01 | PLP | none | m | 57 | 100 | 15 | 27 | right | trauma | none | yes | yes | 0.67 | 1.00 |
| A02 | PLP | HC12 | m | 48 | 75 | 70 | 18 | left | trauma | cosmetic | yes | yes | 3.00 | 3.00 |
| A03 | PLP | HC01 | m | 69 | 57 | 45 | 45 | left | trauma | myo-el. | yes | yes | 3.33 | 2.67 |
| A04 | PLP | HC06 | m | 42 | 85 | 35 | 23 | right | trauma | myo-el. | yes | yes | 0.67 | 1.67 |
| A05 | PLP | HC02 | m | 50 | 73 | 20 | 21 | right | trauma | cosmetic | yes | yes | 4.00 | 0.33 |
| A06 | PLP | HC09 | m | 52 | 85 | 0 | 42 | right | trauma | cosmetic | yes | no | 2.33 | 0.00 |
| A07 | PLP | HC08 | m | 64 | 67 | 0 | 55 | left | tumor | none | yes | yes | 2.00 | 0.00 |
| A08 | PLP | none | m | 62 | 100 | 15 | 43 | right | trauma | none | yes | yes | 2.33 | 2.00 |
| A09 | PLP | none | m | 39 | 80 | 35 | 15 | left | trauma | none | yes | no | 0.33 | 0.00 |
| A10 | PLP | HC15 | m | 37 | 50 | 50 | 13 | left | trauma | cosmetic | yes | yes | 0.33 | 0.00 |
| A11 | PLP | none | m | 30 | 75 | 25 | 19 | right | trauma | none | yes | yes | 2.00 | 2.00 |
| A12 | PLP | none | m | 67 | 83 | 45 | 38 | left | trauma | cosmetic | yes | no | 1.67 | 4.67 |
| A13 | PLP | HC14 | m | 54 | 85 | 21 | 42 | right | trauma | none | yes | yes | 1.67 | 1.33 |
| A14 | PLP | none | m | 73 | 85 | 15 | 15 | right | trauma | none | yes | no | 0.67 | 0.00 |
| A15 | PLP | HC13 | m | 26 | 57 | 38 | 17 | right | trauma | none | yes | yes | 2.33 | 2.00 |
| A16 | PLP | HC04 | m | 42 | 100 | 43 | 19 | left | trauma | none | yes | no | 1.00 | 1.67 |
| A17 | PLP | none | m | 54 | 47 | 15 | 14 | right | trauma | none | yes | yes | 0.67 | 1.00 |
| A18 | PLP | none | m | 63 | 75 | 30 | 40 | right | trauma | myo-el. | yes | yes | 1.67 | 1.00 |
| A19 | PLP | none | m | 52 | 69 | 50 | 16 | right | trauma | manual | yes | no | 1.33 | 2.00 |
| A20 | PLP | none | m | 48 | 76 | 30 | 18 | right | trauma | none | yes | yes | 4.33 | 3.00 |
| A21 | PLP | HC19 | f | 40 | 100 | 45 | 25 | left | other | none | yes | yes | 2.67 | 0.00 |
| A22 | PLP | HC17 | f | 58 | 100 | 25 | 37 | left | trauma | none | yes | yes | 2.33 | 0.67 |
| A23 | PLP | none | f | 53 | 67 | 70 | 23 | right | trauma | none | yes | no | 3.67 | 2.67 |
| A24 | nonPLP | none | m | 59 | 85 | 20 | 10 | right | trauma | myo-el. | no | no | 0.00 | 0.00 |
| A25 | nonPLP | HC05 | m | 67 | 90 | 62 | 19 | right | trauma | myo-el. | yes | yes | 0.00 | 0.00 |
| A26 | nonPLP | HC07 | m | 54 | 100 | 72 | 23 | right | trauma | none | yes | no | 0.00 | 0.00 |
| A27 | nonPLP | none | m | 24 | 83 | 45 | 6 | left | trauma | myo-el. | no | no | 0.00 | missing |
| A28 | nonPLP | none | m | 60 | 57 | 60 | 5 | left | trauma | none | no | no | 0.00 | 0.00 |
| A29 | nonPLP | HC11 | m | 62 | 71 | 10 | 10 | left | trauma | none | no | no | 0.00 | 0.00 |
| A30 | nonPLP | HC10 | m | 47 | 60 | 0 | 22 | left | trauma | none | yes | no | 0.00 | 0.00 |
| A31 | nonPLP | none | m | 48 | 70 | 62 | 45 | right | trauma | none | no | no | 0.00 | 0.00 |
| A32 | nonPLP | HC03 | m | 65 | 69 | 52 | 15 | right | trauma | cosmetic | yes | yes | 0.00 | 0.00 |
| A33 | nonPLP | none | m | 51 | 83 | 10 | 23 | left | trauma | cosmetic | no | no | 0.00 | 0.00 |
| A34 | nonPLP | HC16 | m | 56 | 100 | 65 | 38 | right | trauma | cosmetic | no | no | 0.00 | 0.00 |
| A35 | nonPLP | none | m | 47 | 100 | 25 | 15 | left | tumor | myo-el. | yes | yes | 0.00 | 0.00 |
| A36 | nonPLP | HC18 | f | 52 | 100 | 50 | 18 | left | trauma | myo-el. | no | no | 0.00 | 3.00 |
| A37 | nonPLP | none | f | 51 | 87 | 30 | 28 | left | trauma | none | yes | yes | 0.00 | 0.67 |
| HC01 | HC | A03 | m | 71 | 100 | 45 |  | left |  |  |  |  |  |  |
| HC02 | HC | A05 | m | 50 | 61 | 20 |  | right |  |  |  |  |  |  |
| HC03 | HC | A32 | m | 64 | 100 | 52 |  | right |  |  |  |  |  |  |
| HC04 | HC | A16 | m | 41 | 70 | 43 |  | left |  |  |  |  |  |  |
| HC05 | HC | A25 | m | 67 | 89 | 62 |  | right |  |  |  |  |  |  |
| HC06 | HC | A04 | m | 44 | 75 | 35 |  | right |  |  |  |  |  |  |
| HC07 | HC | A26 | m | 54 | 74 | 72 |  | right |  |  |  |  |  |  |
| HC08 | HC | A07 | m | 64 | 93 | 0 |  | left |  |  |  |  |  |  |
| HC09 | HC | A06 | m | 53 | 89 | 0 |  | right |  |  |  |  |  |  |
| HC10 | HC | A30 | m | 47 | 52 | 0 |  | left |  |  |  |  |  |  |
| HC11 | HC | A29 | m | 63 | 89 | 10 |  | left |  |  |  |  |  |  |
| HC12 | HC | A02 | m | 47 | 64 | 70 |  | left |  |  |  |  |  |  |
| HC13 | HC | A15 | m | 25 | 54 | 38 |  | right |  |  |  |  |  |  |
| HC14 | HC | A13 | m | 51 | 65 | 21 |  | right |  |  |  |  |  |  |
| HC15 | HC | A10 | m | 35 | 89 | 50 |  | left |  |  |  |  |  |  |
| HC16 | HC | A34 | m | 55 | 71 | 65 |  | right |  |  |  |  |  |  |
| HC17 | HC | A22 | f | 58 | 100 | 25 |  | left |  |  |  |  |  |  |
| HC18 | HC | A36 | f | 53 | 31 | 50 |  | left |  |  |  |  |  |  |
| HC19 | HC | A21 | f | 39 | 83 | 45 |  | left |  |  |  |  |  |  |

### 2.2. Body sites

Sensory testing was performed at five body sites. Amputees were tested at the thenar of their intact hand, both corners of the mouth, and the residual limb 5 cm from the distal end of the stump^26,30^ and outside of scar tissue and at a homologous site on the intact arm. Both sites on the arms were marked using cosmetic color and a photo was taken for documentation. The sites on the arms used in the HCs were anatomically homologous to the ones used in the matched amputees.

Body sites for sensory testing were selected based on the following rationale: the residual limb was tested as the body site directly affected by the amputation with a homologous site on the intact arm as a control site. The corners of the mouth were tested because they are remote from the amputation site and clearly outside the area of potential nerve damage. In addition, the face is an interesting area with regard to central changes, since the hand and the face are represented adjacently within the cortical body map in SI. Previous studies showed that an “invasion” (in terms of functional SI reorganization) of the mouth area in the hemisphere contralateral to the amputation into the neighboring former hand area is correlated with the presence of PLP^16,42,45^. It is, however, unclear whether cortical reorganization is reflected in altered sensitivity to stimuli presented at the corners of the mouth. Additionally, we tested the thenar of the existing hand, as sensory changes at the thenar might indicate altered interhemispheric connectivity in the brain because the cortical representation of the existing hand is functionally^29^ and structurally^5^ coupled to the corresponding area in the contralateral hemisphere. Another reason for including the thenar is that it is a standard site for QST^59^.

### 2.3. Sensory testing

Measures were chosen to detect increased or decreased sensitivity to noxious or non-noxious thermal or mechanical stimuli. In total, we assessed four different measures: two pain thresholds—heat pain threshold (HPT) and pressure pain threshold (PPT)—and two detection thresholds—warmth detection threshold (WDT) and tactile two-point discrimination thresholds (2PDT). This allowed us to separate the effects related to the nociceptive system from other somatosensory modalities. The procedures for HPT, WDT and PPT were based on the study protocol developed by the German Research Network on Neuropathic Pain^59^ and were carried out by experimenters trained in QST.

#### 2.3.1. Thermal thresholds

WDT and HPT were measured using a Medoc ATS thermode (3×3 cm surface) included in the Medoc Pathway system (PATHWAY Pain & Sensory Evaluation System, Medoc Ltd. Advanced Medical System, Ramat Yishai, Israel). Ascending (ramp) stimuli were applied with a baseline temperature of 32 C° and a rise rate of 1.5 C°/s (ascending method of limits)^3^. The system’s maximum temperature was set to 52 C° to avoid skin injuries.

When amputees were tested at the thenar and could therefore not respond by button presses, responses, including the ones in the HC, were given verbally to ensure comparability between sites and groups. The assessment of WDT and HPT only differed with respect to the instructions (‘say “stop” as soon as you feel that the temperature is getting warmer’ for WDT and ‘say “stop” when the temperature just becomes painful’ for HPT). When participants said “stop”, the experimenter immediately terminated the ascending stimulus and the thermode returned to the baseline temperature at a rate of 8 C°/s. The participants were asked to keep their eyes closed throughout the test.

In each series, we conducted five consecutive trials. The first trial of each series was presented to familiarize participants with the test and the remaining four stimuli were averaged to calculate the threshold^3^.

#### 2.3.2. Pressure pain thresholds

PPT were measured using a Medoc Algomed (Medoc Ltd. Advanced Medical System, Ramat Yishai, Israel) computerized pressure algometer (surface of 1 cm^2^). Four consecutive ramps were applied using a rate of 50 kPa/s (ascending method of limits)^59^. The participants were instructed to say “stop” when the pressure started to feel painful. Four consecutive stimuli were applied; the first stimulus was excluded and the remaining three were averaged for calculation of the pain threshold.

#### 2.3.3. Two-point discrimination thresholds

2PDT thresholds were measured using a set of 28 calibrated compasses (Rotring “Centro”, Rotring, Hamburg, Germany) ranging in size from 1 to 82 mm in steps of 3 mm. The tips of the compasses were exchanged with blunt rods in order to induce a light touch. If possible, stimuli were applied by placing the compass perpendicularly onto the skin from above. The skin was touched briefly (1–2 s) while the participants kept their eyes closed. We used a simple adaptive staircase (“up-down”) procedure^13,40^, stopping after 7 reversals. The first reversal point was excluded and the following six were averaged to calculate the absolute (50%) threshold. To avoid biases, we used different starting sizes depending on the body site: 25 mm for the corners of the mouth and 46 mm for the arms. 2PDT at the arms were always measured in proximal-to-distal direction while the participant sat in a chair. 2PDT at the corners of the mouth were measured parallel to the inferior-to-superior axis while the participant was lying on a bench to facilitate applying the stimuli perpendicularly from above. Beforehand, the protocol had been evaluated in a pilot study, testing 10 participants’ dorsal forearms. The procedure showed good within-session reliability (Pearson’s *r* = 0.81). The data of this pilot are provided in the supplementary material (see Figure S4). 2PDT was measured in a subsample of 29 amputees and all HC.

### 2.5. Other variables

In order to test whether associations between PLP and sensory thresholds were specific and not mediated via other variables known to be correlates of PLP or related brain plasticity, we also correlated the thresholds to the following variables: use of prostheses^43,11^, (2) embodiment of the prosthesis^33^ (whether the prosthesis is perceived or not perceived as part of the body), telescoping^18,25^, compensation in daily activities by the existing hand^48^, level of the amputation, and RLP^33^. As in a previous study^43^, we calculated an index of prosthesis use by multiplying the type of prosthesis (1=none, 2=cosmetic, 3=myoelectric), the duration for that the prosthesis has been in use, and the frequency of use (per week). Variables were z-transformed before multiplication. The degree of telescoping as well as type, duration and frequency of prosthesis use were assessed using a standardized interview^69,16^. Embodiment of the prosthesis was assessed using a numerical rating scale ranging from 0 („feels like a foreign object“) to 10 („feels like part of my body“) included in the nation-wide survey^4^. Compensatory hand use was assessed with an additional questionnaire^16,43^. Level of amputation was assessed on site and coded as percentage of the length of the intact arm measured from the caput humeri to the tip of the third digit. RLP was assessed in the same way as PLP intensity, using an adaptation of the MPI^16,19,34^ referring to RLP. As for PLP intensity, we used the MPI „pain severity” score. Additionally, we correlated the sensory thresholds with time since amputation and all variables mentioned above were correlated with PLP intensity.

### 2.6. Statistical analyses

#### 2.6.1. Removal of statistical outliers

Prior to statistical analyses, we screened the data for statistical outliers. We used Tukey’s criterion which classifies values as “extreme” outliers if they exceed a range of the third quartile plus 3 interquartile ranges (IQR) or the first quartile minus 3 IQR^62^. To avoid bias in the calculation of the thresholds, we first screened the data of HPT, PPT and WDT at single trial level. We found outliers in 5% of the trial series for WDT and removed single trials. In all other sensory measures the ratios were lower. In WDT and PPT we also removed 2 entire series because less than three values remained for calculation of the thresholds. The number of corrected trial series was comparable between the groups (χ^2^_(2, *N*=56)_ = 0.46, *p* = 0.80).

We also screened the distributions at the threshold level (averages of the single trials) and used the same exclusion criterion as described above. There were no outliers in HPT. In PPT, we removed 2 thresholds (0.7%); in WDT, we removed 7 thresholds (2.5%). The number of removed thresholds was lower in the HC (n = 0) compared to the PLP (n = 4) and nonPLP (n = 5) group (χ^2^_(2, *N*=56)_ = 7.3, *p* = 0.03). A detailed description of the distributions of the raw data and removed outliers is shown in the supplementary material (see Figure S3 and S4).

#### 2.6.2. Statistical models for comparisons between groups and body sites

To compare thresholds between the groups and body sites, we fitted a (repeated measures) linear mixed model (LMM) for each sensory measure using group (PLP, nonPLP and HC) as a between-subject fixed factor and body site (arm ipsilateral, arm contralateral, corner of the mouth ipsilateral, corner of the mouth contralateral, and thenar contralateral to the amputation) as a within-subject fixed factor. LMMs are a more flexible and robust alternative to repeated-measures analyses of variance, and permit simultaneous modeling of discrete grouping variables, repeated factors and continuous covariates^56^. To test whether differences between groups depended on the body site (or vice versa), we also included the interaction between group and body site in the models.

The coding of body site (“ipsilateral” and “contralateral”) in the HC was defined by the side of amputation of the matched amputee. For example, if a HC was matched to a right-sided amputee, his or her right corner of the mouth was coded as “ipsilateral”. Note that in the amputees “ipsilateral arm” refers to the residual limb. This allowed a balanced mixture of right and left side body parts within the categories.

In case of significant effects for either group or body site, we used pairwise post-hoc comparisons to test which variables differed significantly. To avoid alpha error inflation, post-hoc test *p*-values were corrected using the false discovery rate.

#### 2.6.3. Correlational analyses and multiple regression

To test the association between the sensory thresholds and PLP intensity and other continuous variables (described in section 2.4), we calculated Pearson’s correlation coefficients. We correlated each sensory threshold in the PLP group with the “phantom pain severity” score from the MPI. To test whether PLP intensity was associated with a combination of sensory thresholds, we also performed a stepwise multiple regression analysis. At each of the five body sites, we predicted PLP intensity in the PLP group by a combination of HPT, PPT and WDT and determined the best fitting model (based on Akaike’s information criterion). As missing values are a problem for multiple regression approaches, we we imputed missing data (1 value (0.8 %) for HPT and PPT, and 5 values (4.3%) for WDT) by the mean values before the multiple regression analyses. As 2PDT were only assessed in a subset of the participants, this would have resulted in imputation of 18% of the values. Therefore, we excluded 2PDT from the multiple regression analyses.

To test whether correlations between PLP and sensory thresholds were also present across the PLP and nonPLP group together, we also computed correlations between the MPI “pain severity” scores after combining the PLP and nonPLP groups. This indicates whether the sensory theresholds are related to the MPI scores, irrespective of any grouping criterion.

We used the R statistical software^57^. Linear mixed models were performed using the “lme4”^2^ and the “lmerTest”^37^ packages for R. Degrees of freedom in the linear mixed models were estimated using Satterthwaite’s approximation as implemented in the “lmerTest”^37^ package. Figures were created using the “ggplot2” package for R^66^.

## 3. Results

### 3.1. Effects related to the presence of PLP: differences between the groups and body sites

Results from the LMMs for all measures are shown in Fig. 1. Table 2 summarizes the coefficients for the full models (including the factor group and body site, and the group * body site interaction). Table 3 summarizes the results from post-hoc comparisons between the body sites.

**Figure 1:**
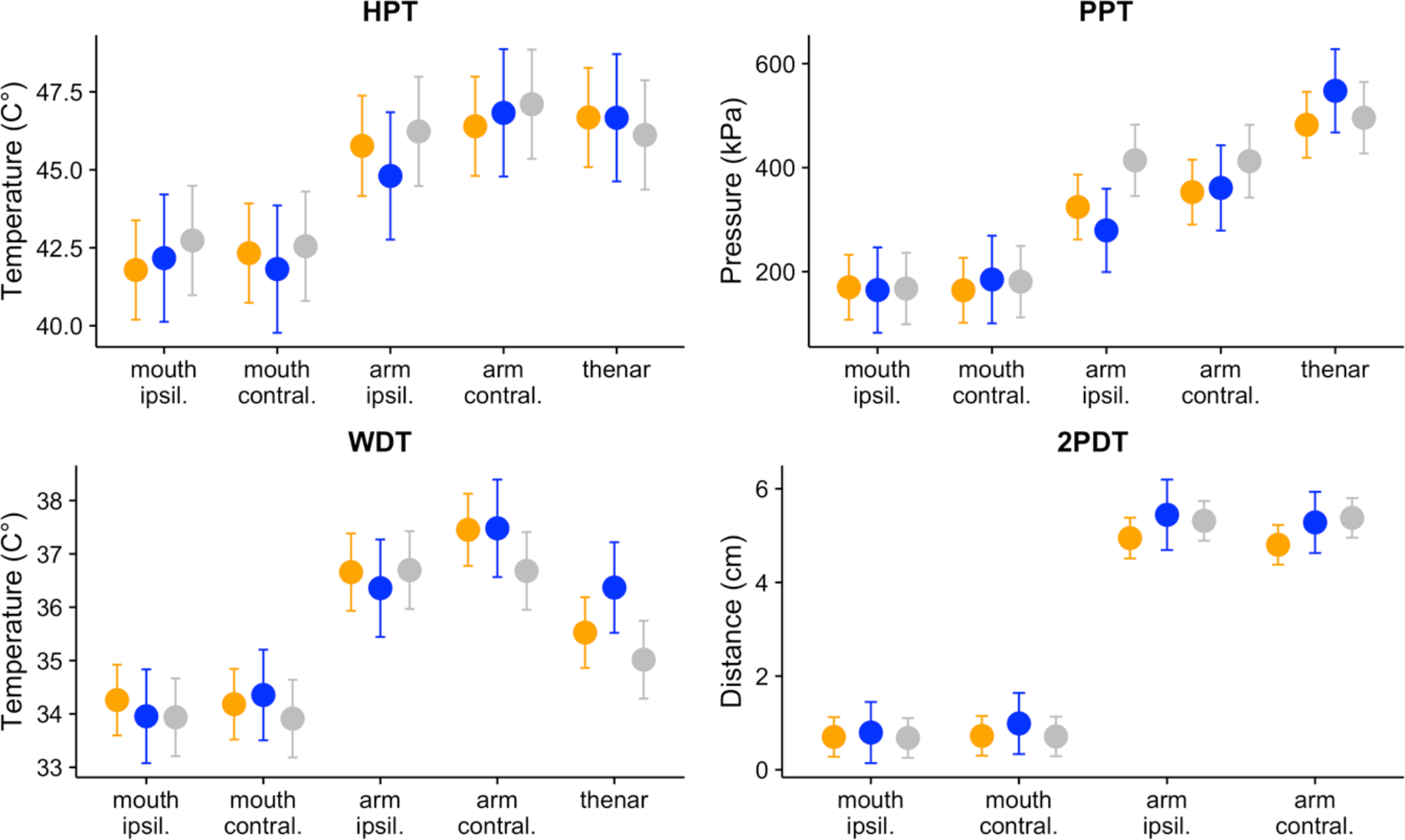
Results of the linear mixed models comparing the thresholds for heat pain (HPT), pressure pain (PPT), warmth detection (WDT) and two-point discrimination (2PDT) between the groups (amputees with phantom limb pain: orange, amputees without phantom limb pain: blue; healthy controls: grey) and the body sites. The values represent the models’ effects expressed as least squares means (fitted values) with 95% confidence intervals (shown as error bars). There were no significant differences between the groups at any of the comparisons. The body sites differed significantly (see the text as well as Table 2 and 3 for details).

**Table 2:** Results for main effects and interactions from linear mixed models for heat pain thresholds (HPT), pressure pain thresholds (PPT), warmth detection thresholds (WDT) and two-point discrimination thresholds (2PDT) including the factors group, body site, and their interaction.

| Measure | Term | F | DF (num./denom.) | P |
| --- | --- | --- | --- | --- |
| HPT | Group | 0.10 | 2/53 | 0.901 |
|  | Body site | 50.48 | 4/211 | < 0.001 |
|  | Group * Body site | 0.65 | 8/211 | 0.732 |
| PPT | Group | 0.51 | 2/53 | 0.605 |
|  | Body site | 87.12 | 4/207 | < 0.001 |
|  | Group * Body site | 1.80 | 8/207 | 0.079 |
| WDT | Group | 1.23 | 2/53 | 0.3 |
|  | Body site | 45.52 | 4/204 | < 0.001 |
|  | Group * Body site | 0.88 | 8/204 | 0.534 |
| 2PDT | Group | 1.43 | 2/44 | 0.25 |
|  | Body site | 304.21 | 3/129 | < 0.001 |
|  | Group * Body site | 0.54 | 6/128 | 0.775 |

**Table 3:** Results from post-hoc comparisons for the significant factor body site. Post-hoc tests followed linear mixed models for each of the sensory thresholds (heat pain thresholds (HPT), pressure pain thresholds (PPT), warmth detection thresholds (WDT) and two-point discrimination thresholds (2PDT. Within each model, p values were adjusted for multiple comparisons using false discovery rate.

| Measure | Comparison | Estimate | SE | z | P |
| --- | --- | --- | --- | --- | --- |
| HPT | arm ipsil. - mouth ipsil. | 3.98 | 0.70 | 5.70 | < 0.001 |
|  | thenar - mouth ipsil. | 4.89 | 0.69 | 7.10 | < 0.001 |
|  | mouth contral. - mouth ipsil. | 0.54 | 0.69 | 0.78 | 0.481 |
|  | arm contral. - mouth ipsil. | 4.61 | 0.69 | 6.70 | < 0.001 |
|  | thenar - arm ipsil. | 0.91 | 0.70 | 1.30 | 0.276 |
|  | mouth contral. - arm ipsil. | -3.44 | 0.70 | -4.93 | < 0.001 |
|  | arm contral. - arm ipsil. | 0.63 | 0.70 | 0.90 | 0.461 |
|  | mouth contral. - thenar | -4.35 | 0.69 | -6.32 | < 0.001 |
|  | arm contral. - thenar | -0.28 | 0.69 | -0.41 | 0.683 |
|  | arm contral. - mouth contral. | 4.07 | 0.69 | 5.91 | < 0.001 |
| PPT | arm ipsil. - mouth ipsil. | 154.28 | 32.60 | 4.73 | < 0.001 |
|  | thenar - mouth ipsil. | 312.37 | 33.04 | 9.45 | < 0.001 |
|  | mouth contral. - mouth ipsil. | -5.83 | 32.60 | -0.18 | 0.858 |
|  | arm contral. - mouth ipsil. | 182.95 | 32.60 | 5.61 | < 0.001 |
|  | thenar - arm ipsil. | 158.09 | 33.04 | 4.79 | < 0.001 |
|  | mouth contral. - arm ipsil. | -160.11 | 32.60 | -4.91 | < 0.001 |
|  | arm contral. - arm ipsil. | 28.68 | 32.60 | 0.88 | 0.421 |
|  | mouth contral. - thenar | -318.20 | 33.04 | -9.63 | < 0.001 |
|  | arm contral. - thenar | -129.42 | 33.04 | -3.92 | < 0.001 |
|  | arm contral. - mouth contral. | 188.78 | 32.60 | 5.79 | < 0.001 |
| WDT | arm ipsil. - mouth ipsil. | 2.40 | 0.46 | 5.17 | < 0.001 |
|  | thenar - mouth ipsil. | 1.27 | 0.44 | 2.88 | 0.006 |
|  | mouth contral. - mouth ipsil. | -0.08 | 0.44 | -0.18 | 0.86 |
|  | arm contral. - mouth ipsil. | 3.19 | 0.44 | 7.18 | < 0.001 |
|  | thenar - arm ipsil. | -1.13 | 0.46 | -2.44 | 0.018 |
|  | mouth contral. - arm ipsil. | -2.48 | 0.46 | -5.34 | < 0.001 |
|  | arm contral. - arm ipsil. | 0.79 | 0.47 | 1.70 | 0.1 |
|  | mouth contral. - thenar | -1.34 | 0.44 | -3.06 | 0.004 |
|  | arm contral. - thenar | 1.93 | 0.44 | 4.33 | < 0.001 |
|  | arm contral. - mouth contral. | 3.27 | 0.44 | 7.35 | < 0.001 |
| 2PDT | mouth contral. - mouth ipsil. | 0.02 | 0.29 | 0.08 | 0.937 |
|  | arm ipsil. - mouth ipsil. | 4.25 | 0.29 | 14.41 | < 0.001 |
|  | arm contral. - mouth ipsil. | 4.10 | 0.29 | 14.13 | < 0.001 |
|  | arm ipsil. - mouth contral. | 4.22 | 0.29 | 14.33 | < 0.001 |
|  | arm contral. - mouth contral. | 4.08 | 0.29 | 14.05 | < 0.001 |
|  | arm contral. - arm ipsil. | -0.14 | 0.29 | -0.48 | 0.753 |

#### 3.1.1 Heat Pain Thresholds

There was no significant main effect of group (F_(2, 53)_ = 0.1, *p* = 0.901) and no significant interaction between group and body site (F_(8, 211)_ = 0.65, *p* = 0.732). The main effect of body site was significant (F_(4, 211)_ = 50.5, *p* < 0.001). Post-hoc camparisons (see Table 3) showed that thresholds were lower at both corners of the mouth compared to both arms and the thenar. The corners of the mouth did not differ significantly from each other, and the thenar did not differ significantly from both arms. Noteworthy, the thresholds were lower, albeit not statistically significant, at the ipsilateral arm compared to the contralateral arm.

#### 3.1.2 Pressure Pain Thresholds

For PPT, there was no significant effect of group (F_(2, 53)_ = 0.51, *p* = 0.605) or group * body site (F_(8, 207)_ = 1.8, *p* = 0.079). The factor body site was significant (F_(4, 207)_ = 87.1, *p* < 0.001). Post-hoc comparisons between body sites revealed that all body sites differed from each other except for the homologous body parts at the corners of the mouth and the arms, which did not differ significantly.

#### 3.1.3 Warmth Detection Thresholds

No significant effect of group (F_(2, 53)_ = 1.23, *p* = 0.3) and group * body site was found (F_(8, 204)_ = 0.88, *p* = 0.534). The factor body site was significant (F_(4, 204)_ = 45.52, *p* < 0.001). Post-hoc comparisons revealed significant differences between all body sites, except for the comparison between the corners of the mouth. The comparison between both arms was not significant.

#### 3.1.4 Two-Point Discrimination Thresholds

For 2PDT, there was no significant effect of group (F_(2, 44)_ = 1.43, *p* = 0.25) or group * body site (F_(6, 128)_ = 0.54, *p* = 0.775). Body site was significant (F_(3, 129)_ = 304.21, *p* < 0.001). Post-hoc tests showed that thresholds were lower at both corners of the mouth than at both arms. The homologous parts at the corners of the mouth and at the arms did not differ significantly from each other.

### 3.2. Effects related to the intensity of PLP: correlational analyses

#### 3.2.1. Correlations between PLP intensity and sensory thresholds

All correlations between the sensory thresholds in the PLP group and the reported intensity of their PLP are shown in Fig. 2. For HPT, the correlations were negative, indicating that more intense PLP was associated with lower HPT, and significant for the ipsilateral corner of the mouth (r = −0.59, *p* = 0.003), the contralateral corner of the mouth (r = −0.65, *p* < 0.001), the thenar (r = −0.64, *p* < 0.001), and the contralateral arm (r = −0.45, *p* = 0.031. Only at the ipsilateral arm (residual limb), the correlation was not significant (r = −0.33, *p* = 0.137). PPT were significantly negatively correlated with PLP intensity at the contralateral corner of the mouth (r = −0.47, *p* = 0.024). At all other body sites, the correlations were negative, but not significant (see Fig. 2). WDT were not significantly correlated with PLP intensity (see Fig. 2). Numerically, the correlations were highest at the ipsilateral corner of the mouth where the correlation was positive and a statistical trend (r = 0.35, *p* = 0.097). 2PDT were not significantly correlated with PLP intensity at all body sites (see Fig. 2).

**Figure 2:**
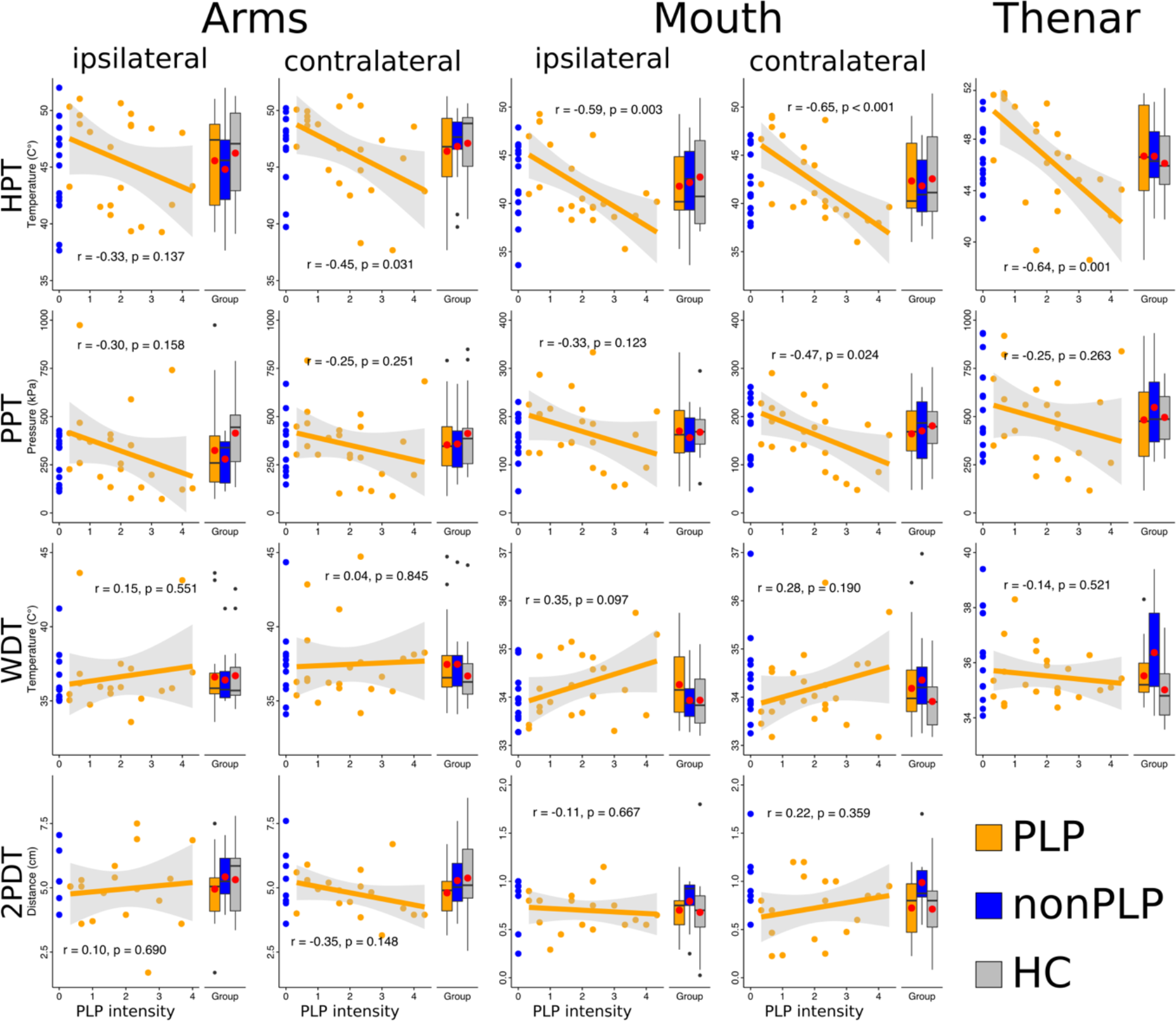
Distributions of somatosensory thresholds and their correlations with the intensity of phantom limb pain (PLP) in the PLP group (orange). The rows represent the thresholds, which were assessed: heat pain thresholds (HPT), pressure pain thresholds (PPT), warmth detection thresholds (WDT) and two-point discrimination thresholds (2PDT). The columns represent the body sites: the arm ipsilateral to the amputation (in amputees the residual limb, in healthy controls a matched site), the contralateral arm, the ipsilateral and the contralateral corner of the mouth, and the contralateral thenar. Within each subplot, the scatter plots (on the left) show the data of the PLP group and the amputees without PLP (nonPLP; blue) for the respective threshold and PLP intensity. For the PLP group, the result of a linear regression model is shown. The gray-shaded area around the regression line indicates the borders of the 95% confidence interval. The Pearson correlation coefficient together with a (two-sided) p-value is shown within the plot. The right side of each subplot shows the distribution of the sensory thresholds as boxplots for the PLP group, the nonPLP group and healthy controls (HC; grey). The mean value for each group is highlighted as a red point.

When correlations were computed after combining the PLP and the nonPLP group, significant negative correlations were found for HPT at both the ipsilateral (r = −0.36, *p* = 0.030) and the contralateral (r = −0.34, *p* = 0.042) corner of the mouth as well as at the thenar (r = −0.39, *p* = 0.016). There was also a significant positive correlation for WDT at the ipsilateral corner of the mouth (r = 0.38, *p* = 0.022). All other correlations were not significant (all p > 0.67).

#### 3.2.2. Correlations between PLP intensity and differences between sensory thresholds at homologous body sites

Intra-individual differences between the homologous areas at the arms and the corners of the mouth were not significantly correlated with PLP intensity for any of the sensory thresholds (all r > −0.32 and < 0.34, all *p* > 0.10).

When the PLP and the nonPLP group were combined, there was a significant correlation for the difference between PPT at the corners of the mouth, indicating that lower PPT at the ipsilateral compared to the contralateral corner of the mouth was associated with lower MPI scores (r = 0.36, *p* = 0.033). All other correlations for the groups combined were not significant (all r > 0.06 and < 0.12, all *p* > 0.14).

All correlations are shown in the supplementary material.

#### 3.2.3. Stepwise multiple regressions per body site

At the ipsilateral arm, the stepwise procedure removed all sensory thresholds except for HPT, which remained as the only predictor in the best fitting model. However, this model was not significant (*F*_(1, 22)_ = 2.34, *p* = 0.14). At the contralateral thenar the best fitting model also only included HPT. This model was significant and explained 41% of the variance in PLP intensity (*F*_(1, 21)_ = 14.45, *p* = 0.001; R^2^ = 0.41; R^2^_adj._ = 0.38). At both the ipsilateral and the contralateral corner of the mouth, the best fitting model included both HPT and WDT. HPT received a negative and WDT a positive weight (see Fig. 3), indicating that PLP intensity was associated with lower HPT, i.e., enhanced pain sensitivity, and higher WDT, i.e., reduced non-painful thermal sensitivity. At the ipsilateral corner of the mouth, this model was significant and explained 44% of the variance in PLP intensity (*F*_(2, 20)_ = 7.73, *p* = 0.003; R^2^ = 0.44; R^2^_adj._ = 0.38). The model at the contralateral corner of the mouth was also significant and explained 58% of the variance in PLP intensity (*F*_(2, 20)_ = 14.01, *p* < 0.001; R^2^ = 0.58; R^2^_adj._ = 0.54). The results of the best fitting multiple regression for both corners of the are shown in Fig. 3, the results of the bivariate correlations for HPT at the ipsilateral arm and the contralateral thenar are included in Fig. 2.

**Figure 3:**
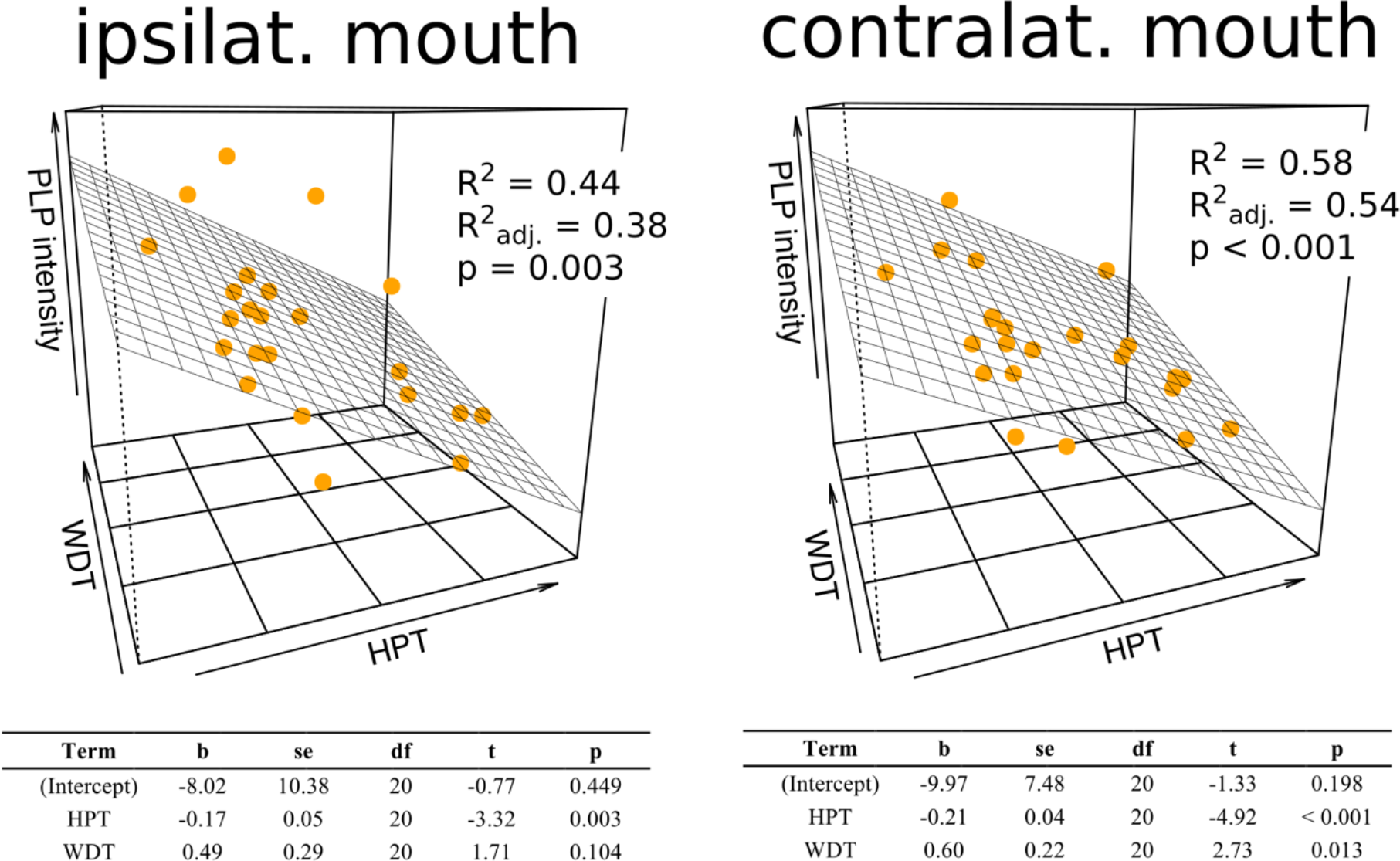
Results of the multiple regression models in which the intensity of phantom limb pain (PLP intensity) in the group of amputees with phantom limb pain was statistically predicted by a linear combination of somatosensory thresholds measured all on the same body site. At the corners of the mouth, both ipsilateral and contralateral to the amputation, PLP intensity was predicted by a combination of heat pain thresholds (HPT) and warmth detection thresholds (WDT). The three-dimensional scatter plots show the data together with a regression plane, representing the values predicted by the multiple regression model. The proportion of variance in PLP intensity explained by the model is shown within the scatter plots. The tables on the bottom right show the regression coefficients for each of the models. At body sites not shown in the figure, the models were not significant (ipsilateral arm) or PLP intensity was only predicted by HPT (contralateral arm and contralateral thenar).

### 3.3. Additional correlational analyses

All correlations are presented in Table 4. Time since amputation (all *p* > 0.05) as well as embodiment of the prosthesis (all *p* > 0.06) were not significantly correlated with any of the sensory thresholds or PLP intensity. A higher degree of prosthesis use was associated with lower HPT at the ipsilateral arm (r = −0.496, *p* < 0.01) and with higher WDT at the contralateral thenar (r = 0.405, *p* = 0.02). The degree of compensatory use of the existing hand was associated with higher WDT at the ipsilateral arm (r = 0.389, *p* = 0.03) and at the contralateral thenar (r = 0.336, *p* = 0.05) as well as higher PPT at the ipsilateral (r = 0.384, *p* = 0.02) and the contralateral corner of the mouth (r = 0.469, *p* = 0.01). The percentage of telescoping (all *p* > 0.24) and RLP (all *p* > 0.10) were both not significantly correlated to any of the sensory thresholds. However, both telescoping (r = 0.392, *p* = 0.02) and RLP (r = 0.518, *p* < 0.01) were significantly associated with higher PLP intensity. There were significant negative correlations of a medium size between the level of amputation with HPT (r = −0.527, *p* < 0.01) and WDT (r = −0.518, *p* < 0.01) at the ipsilateral arm indicating lower thresholds for more distal amputations.

As HPT at the ipsilateral arm was significantly correlated with level of amputation and with degree of prosthesis use, we performed an additional multiple regression analysis including both factors to clarify whether they explain the same or different proportions of the variance in HPT at the residual limb. In the multiple regression model, both level of amputation (*b* = −0.080, t_31_ = −2.68, *p* = 0.01) and degree of prosthesis use (*b* = −0.312, t_31_ = −2.63, *p* = 0.01) were significant and were therefore independent predictors of HPT.

**Table 4:**
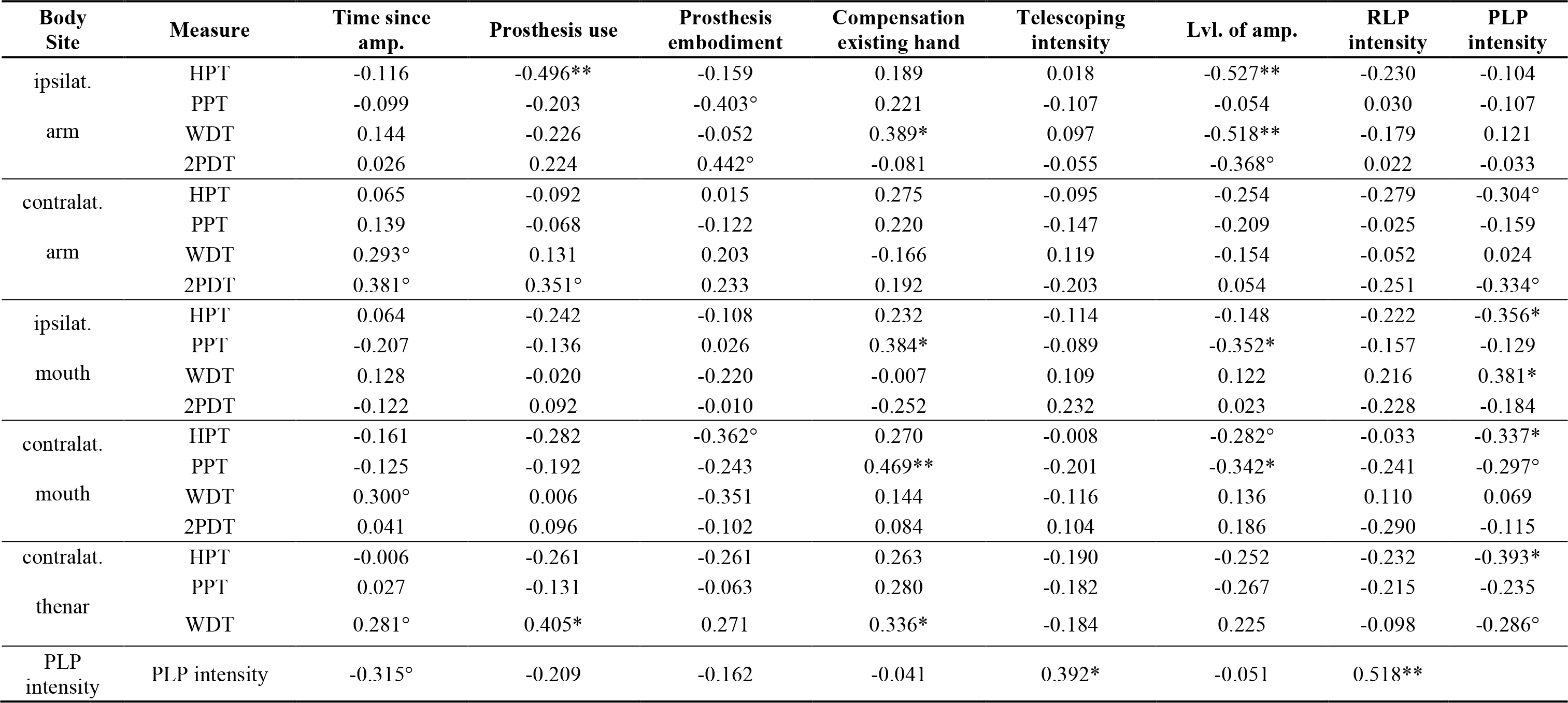
Pearson correlations between heat pain thresholds (HPT), pressure pain thresholds (PPT), warmth detection thresholds (WDT) and two-point discrimination thresholds (2PDT) at the measured body sites and time since amputation, prosthesis use, perceived embodiment of the prosthesis, the degree of compensation with the existing hand, and the intensity of telescoping, level of amputation, residual limb pain (RLP) and phantom limb pain (PLP). The correlations are based on both amputee groups together (with PLP: n=23; without PLP: n=14). *p<0.05, **p<0.01.

## 4. Discussion

This study investigated whether changes in nociceptive and non-nociceptive somatosensory thresholds assessed at different body sites correlate with the presence and/or the perceived intensity of PLP in upper limb amputees. We tested for associations with the presence of PLP by comparing pain and perception thresholds measured in the PLP group to those measured in the nonPLP group and in HC. Surprisingly, there were no significant differences between HPT and PPT at any of the measured body sites as a function of *absence/presence* of PLP. However, we found strong relationships between pain thresholds and the *intensity* of PLP in the PLP group. Higher PLP intensity was associated with lower HPT at all body sites except for the residual limb. Positive correlations were also found for PPT, however, the relationships were weaker and significant only at the contralateral corner of the mouth. For a comparison, we also computed correlations between PLP intensity across the PLP and the nonPLP group together. Especially for HPT, associations remained significant, although numerically lower. While an association of HPT with the presence of PLP cannot be completely ruled out based on our data, we suggest that it is mainly the intensity and not the presence of PLP that is associated with pain sensitivity.

We did not find evidence for an association between perception thresholds and the presence of PLP, as no significant group differences were observed. Differently from pain thresholds, there were no significant associations with the intensity of PLP in the PLP group either. However, in contrast to the pain thresholds, for which the correlations were negative, correlations for WDT were numerically positive at all body sites except for the thenar. Taken together, higher PLP intensity correlated with lower HPT (enhanced pain sensitivity) but with higher WDT (reduced non-nociceptive thermal sensitivity). This differential relationship of PLP intensity with HPT versus WDT was also apparent from the results of stepwise multiple regressions run within the PLP group. At the corners of the mouth, the best fitting models included both HPT and WDT, which together explained a very high proportion (of up to 58%) of the variance in PLP intensity. Correlations between the sensory thresholds were also specifically associated with PLP. By way of contrast, RLP, frequently reported in amputees with PLP^33^ and also positively correlated with PLP in this study, was not significantly associated with any of the sensory thresholds.

HPT were significantly lower at the residual limb compared to the intact limb, which could be viewed as a sign of peripheral hyperalgesia. Importantly, this local hyperalgesia was observed in both the PLP and in the nonPLP group. Moreover, the difference between the limbs was also not significantly correlated with PLP intensity. Correlational analyses showed instead that HPT at the ipsilateral arm significantly correlated with the use of prostheses. A higher degree of prosthesis use was associated with lower HPT, which could be due to the prosthesis inducing irritation or compression of the residual limb^6,12,44^. We also found that HPT at the residual limb was significantly lower for longer residual limbs, when sensory testing was carried out in more distal parts of the limb. Similar proximal-to-distal gradients have already been shown for tactile sensitivity in the residual limbs of amputees^27^. Using multiple regression analyses, we also demonstrated that both these factors—more prosthesis use and higher level of amputation—were independently related to lower HPT. We revisited the hypothesis of increased tactile acuity of the residual limb. Contrary to early^27,31,61,68^, but in line with more recent studies^17,25,30^, we did not find significantly lower 2PDT at the residual limb. There were also no significant differences between the limbs in WDT. However, similar to what we found for HPT, WDT at the residual limb were significantly lower for increased use of prostheses. Taken together, these results support the conclusion drawn by Hunter et al.^30^ that sensory measures taken at the residual limb do not in any simple way reflect phantom phenomena. Our data further suggest that they are instead more strongly related to the use of prostheses and RLP.

Because cortical reorganization is both associated with PLP^16^ and with sensory changes in the face in healthy subjects^51^, sensory thresholds might be altered in amputees’ corners of the mouth. However, we did neither find significant differences between the groups at the the corners of the mouth nor were there significant intraindividual differences between the corners of the mouth for any of the thresholds. This indicates that the functional reorganization in S1 is not necessarily related to basic perceptional changes, suggesting that other central mechanisms are involved.

In summary, our measurements of pain thresholds at different body sites, remote from the residual limb, and their interrelationships, suggest that global enhanced pain sensitivity is related to PLP. We suggest that this global sensitivity reflects a central aspect of sensory processing rather than peripheral mechanisms and that it is evidence for the presence of “widespread hypersensitivity” already suggested by Cronholm^7^ in 1951 (p. 119). Whether this phenomenon relates to plasticity processes occurring after or to factors already present before amputation, cannot be clarified based on the present data. Central sensitization resulting from long-term nociceptive input is a potential explanation for the correlation between PLP intensity and enhanced pain sensitivity^72^ and the generalization of pain sensitivity to remote areas of the body^39^ in participants with strong PLP. However, it is also possible that participants who developed strong PLP already had low pain thresholds prior to amputation^53^.

In any case, it seems unlikely that low pain thresholds alone may be a risk factor for developing PLP or a necessary consequence of the presence of PLP. Low pain thresholds were seen in participants in the PLP group but also in the nonPLP group. Nonetheless, the participants in the nonPLP group did obviously not develop chronic PLP. Conversely, some participants in the PLP group manifested a type of PLP, which did not coincide with low pain thresholds. In fact, participants with low intensity PLP showed high pain thresholds. This illustrates that there might be a dissociation between the role of pain thresholds in the absence versus presence of PLP, such that pain thresholds are not predictive of whether pain is present or not but, if pain is present, predict its intensity. Davis^8^ recently suggested that the presence of pain is ultimately a qualitatively different state from the absence of pain. Factors such as intensity or quality merely modulate the pain experience. As much as it is known that the presence of chronic pain changes neural processing of sensory information^9,38,72^, memory, affective learning and emotional processes^20^, we suggest that pain sensitivity might play a different role when chronic pain is present and interact with the intensity of the chronic pain. However, the factors determining whether amputees develop chronic pain or not are manifold^15^ and altered pain processing might only be one among them^18,53,54^. In the last years, functional^1^ and structural^50^ brain connectivity have been shown to predict the transition from acute to chronic back pain in longitudinal studies. Although PLP patients might differ from chronic back pain patients especially with respect to the role of cognitive and emotional variables^22,23^, it should be clarified whether brain connectivity also predicts chronic PLP and how this relates to sensory processing and cortical reorganization.

There are some limitations to the present study. First, the study is only based on psychophysical thresholds. Therefore, no conclusions can be drawn as to whether the origin of enhanced pain sensitivity is due to peripheral, spinal or cerebral mechanisms. Secondly, the study relied on a cross-sectional design and its findings are only correlational. It is both possible that increased pain sensitivity develops as a consequence of chronic PLP or, alternatively, that high pain sensitivity has been present as a pre-amputation trait, potentially promoting the emergence of PLP. Preoperation pain sensitivity has been shown to predict postoperative pain^24^ and PLP early after amputation correlated with pain sensitivity assessed directly before the amputation^53^. However, both for PLP and for other chronic pain following injury, little is known about whether later-stage chronic pain is also associated with pain sensitivity before injury. As these relevant questions need to be addresses using a longitudinal design, the present study can not contribute to clarify these etiological questions.

## Acknowledgment

This study was supported by a grant from the Deutsche Forschungsgemeinschaft (SFB1158/B07) to HF and JA and European Research Council Advanced grant “Phantom phenomena: A window to the mind and the brain” (PHANTOMMIND, FP7/2007–2013/230249) to HF.

We would also like to thank Kristina Staudt and student assistants for their help with data acquisition.

## Data Availability

Data and code of this study are publicly available and can be accessed via the Open Science Framework (https://osf.io/v6hjy/).

