## Supplementary Material for "Phantom limb pain intensity is associated with generalized hyperalgesia"

*Table of contents*

Sensory data: descriptive table (means and standard deviations for all assessments)

Correlations between intraindividual side differences and PLP intensity

Details about outlier removal

Results from a pilot study assessing the reliability and validity of the two-point discrimination test

### Sensory data: descriptive table

| Measure | Group | ipsilateral arm | | | contralateral arm | | | ipsilateral mouth | | | contralateral mouth | | | contralateral thenar | | |
|---|---|---|---|---|---|---|---|---|---|---|---|---|---|---|---|---|
| | | n | M | SD | n | M | SD | n | M | SD | n | M | SD | n | M | SD |
| HPT (C°) | PLP | 22 | 45.56 | 4.15 | 23 | 46.40 | 3.75 | 23 | 41.79 | 3.96 | 23 | 42.33 | 4.10 | 23 | 46.68 | 3.98 |
| | nonPLP | 14 | 44.80 | 4.20 | 14 | 46.83 | 3.18 | 14 | 42.17 | 4.06 | 14 | 41.82 | 3.25 | 14 | 46.67 | 2.60 |
| | HC | 19 | 46.23 | 4.08 | 19 | 47.10 | 3.15 | 19 | 42.74 | 4.88 | 19 | 42.55 | 4.59 | 19 | 46.12 | 2.82 |
| PPT (kPa) | PLP | 23 | 324.24 | 218.85 | 23 | 352.92 | 178.50 | 23 | 169.96 | 71.51 | 23 | 164.13 | 66.10 | 22 | 482.58 | 226.09 |
| | nonPLP | 14 | 279.37 | 119.00 | 13 | 357.64 | 147.71 | 13 | 156.22 | 50.67 | 12 | 170.18 | 69.64 | 14 | 547.70 | 233.08 |
| | HC | 19 | 414.04 | 176.81 | 18 | 412.36 | 203.33 | 19 | 167.50 | 49.40 | 19 | 180.73 | 58.71 | 19 | 495.88 | 143.54 |
| WDT (C°) | PLP | 19 | 36.61 | 2.55 | 22 | 37.46 | 2.56 | 23 | 34.26 | 0.68 | 23 | 34.18 | 0.78 | 23 | 35.52 | 0.96 |
| | nonPLP | 12 | 36.41 | 1.83 | 12 | 37.47 | 2.62 | 13 | 33.93 | 0.55 | 14 | 34.35 | 0.92 | 14 | 36.37 | 1.64 |
| | HC | 19 | 36.70 | 2.13 | 19 | 36.68 | 2.21 | 19 | 33.94 | 0.58 | 19 | 33.91 | 0.59 | 19 | 35.02 | 1.11 |
| 2PDT (cm) | PLP | 18 | 4.95 | 1.38 | 19 | 4.80 | 0.86 | 19 | 0.70 | 0.21 | 19 | 0.72 | 0.31 | 0 | NA | NA |
| | nonPLP | 6 | 5.42 | 1.15 | 8 | 5.28 | 1.26 | 8 | 0.79 | 0.28 | 8 | 0.99 | 0.34 | 0 | NA | NA |
| | HC | 19 | 5.31 | 1.27 | 19 | 5.38 | 1.60 | 19 | 0.68 | 0.40 | 19 | 0.71 | 0.38 | 0 | NA | NA |

Table S2: Number of valid observations (n), mean values (M) and standard deviations for all sensory thresholds assessed in the study. Values for heat pain thresholds (HPT), pressure pain thresholds (PPT), warmth detection thresholds (WDT) and two-point discrimination thresholds (2PDT) are shown separately for the group with phantom limb pain (PLP), the group without PLP (nonPLP) and the healthy controls (HC).

**Correlations between PLP intensity and differences in sensory thresholds between homologous areas on both sides of the body**

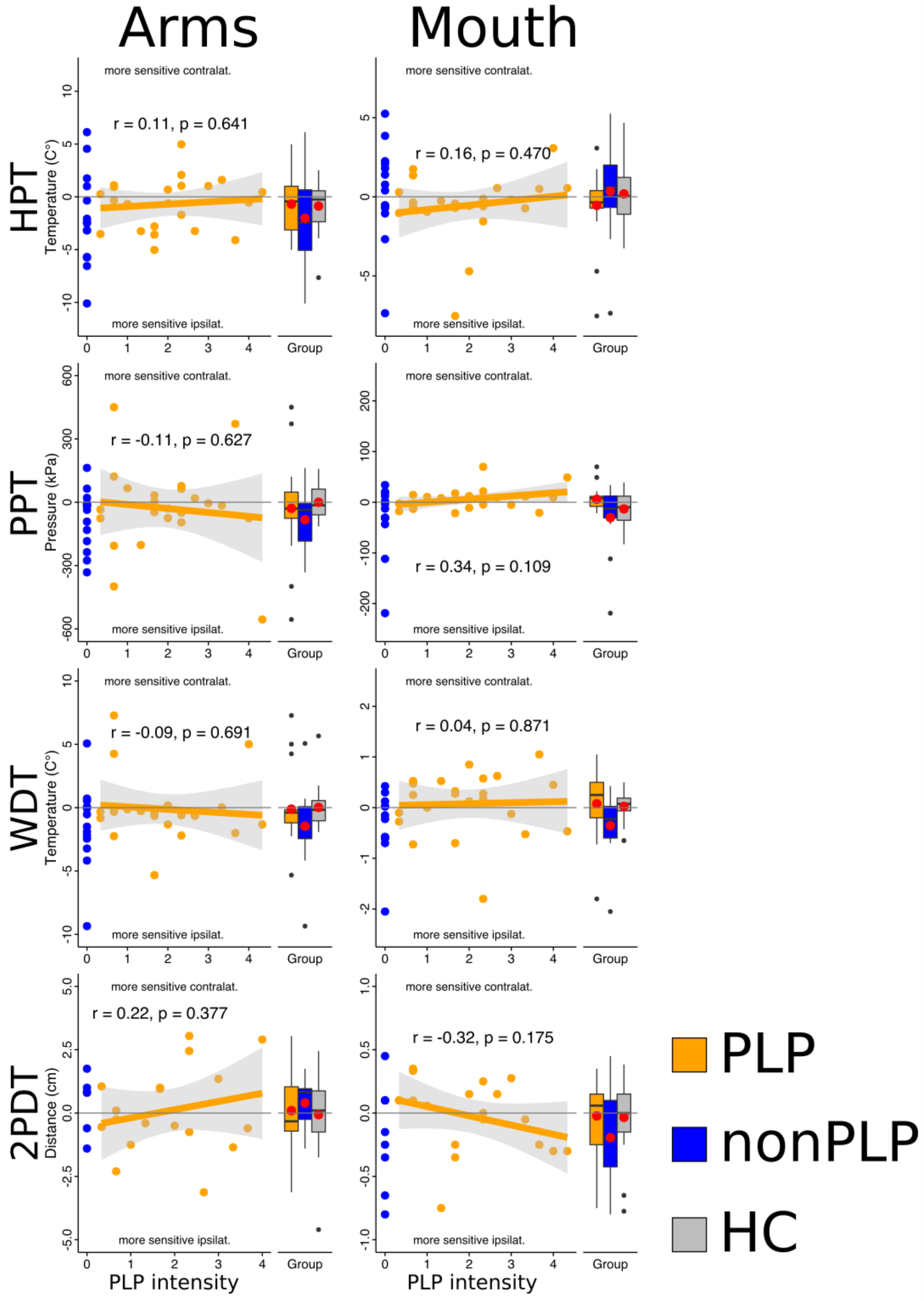


Figure S1: Differences of somatosensory thresholds between homologous areas on both sides of the body (at the arms and the corners of the mouth) and the correlations of theses difference values with the intensity of phantom limb pain (PLP) in the PLP group (orange). The rows represent the thresholds, which were assessed: heat pain thresholds (HPT), pressure pain thresholds (PPT), warmth detection thresholds (WDT) and two-point discrimination thresholds (2PDT). Difference values were computed by subtracting the threshold of the site contralateral from the site ipsilateral to the amputation. The columns represent the pairs of homologous body sites: the arms and the corners of the. Within each subplot, the scatter plots (on the left) show the data of the PLP group and the amputees without PLP (nonPLP; blue) for the respective difference value and PLP intensity. For the PLP group, the result of a linear regression model is shown. The gray-shaded area around the regression line indicates the borders of the 95% confidence interval. The Pearson correlation coefficient together with a (two-sided) p-value is shown within the plot. The right side of each subplot shows the distribution of the difference values as boxplots for the PLP group, the nonPLP group and healthy controls (HC; grey). The mean value for each group is highlighted as a red point.

### Data screening and removal of outliers

Prior to statistical analyses we screened the distributions of the WDT, HPT and PPT and removed statistical outliers. To detect outliers at the single-trial level, we analyzed intra-individual variations over the trials. Within each participant, we standardized the data by subtracting the mean value (over the trials of the participant) from the original values. This resulted in values expressing each trial's deviation from the mean. We then analyzed the distribution of these standardized values for each trial at the level of entire sample (all groups collapsed) to determine if there were outliers. Outliers were defined using Tukey's criterion, which classifies values as outliers if they are either below the first quartile minus 3 interquartile ranges (IQR) or above the third quartile plus 3 IQR. Outliers were removed from the data and the thresholds were calculated by averaging the remaining trials. If, however, more than 50% of the data had been removed, all data were discarded and no threshold was computed.

The distributions of the standardized values and outliers are shown in Figure S1. In both HPT and PPT, outliers were removed in 6 series (2.1%). In PPT two series of trials (0.7%) were removed completely because of more than 50% outliers. In WDT, outliers were removed in 14 (5%) of the series and two series (0.7%) were removed completely.

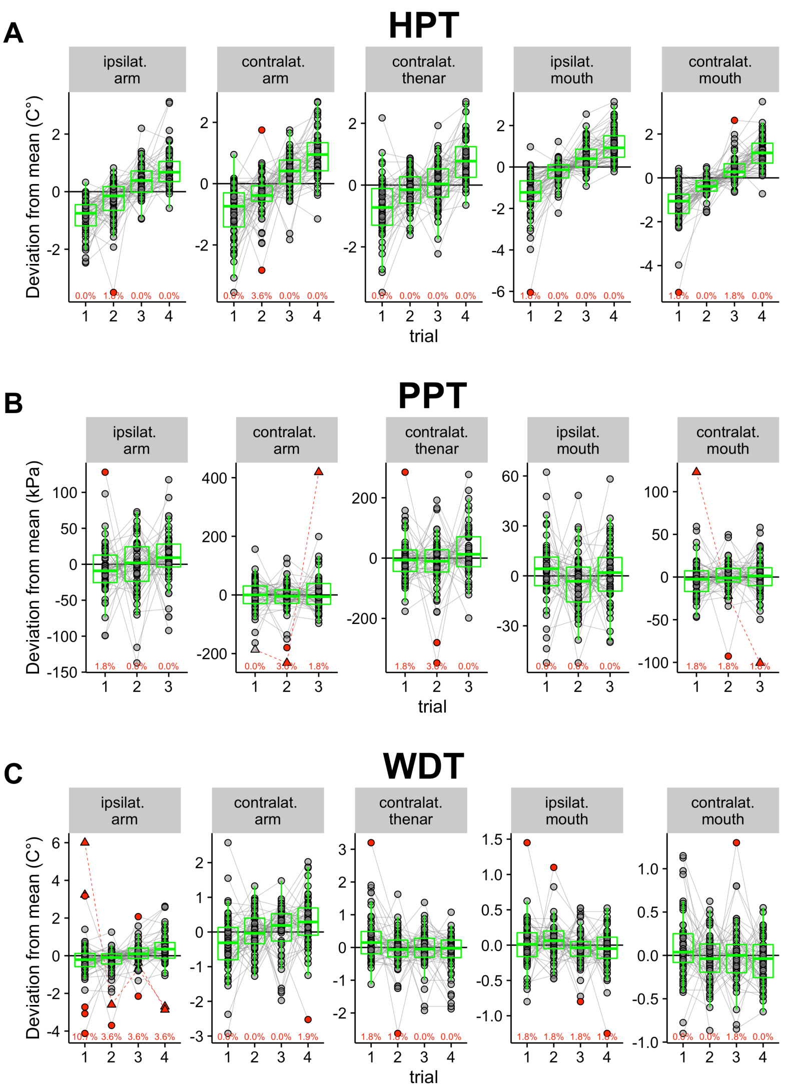


Figure S2: Single-trial data for (A) heat pain thresholds (HPT), (B) pressure pain thresholds (PPT) and (C) warmth detection thresholds (WDT) for all participants. The abscissa represents the trials (4 for HPT and WDT and 3 for PPT) and the ordinate represents the trials' deviation from each participant's mean value (aggregated over the trials). For each trial, the distribution at the group level is shown by green boxplots. Data belonging to the same participant are connected with lines. Gray points range between the third quartile + 3 interquartile ranges (IQR) and the first quartile - 3 IQR. Red symbols exceed this range and are classified as outliers. Red points highlight observations that were removed before calculating the threshold (defined as the average of the valid trials in original units). Triangles that are interconnected with dotted red lines highlight cases in which more than 50% of the values belonging to the participant were classified as outliers and in which case we discarded all trials and did not compute a threshold. The proportion of values classified as outliers (in percent) is shown in red, below the boxplots.

After calculating all thresholds based on the remaining trials for HPT, PPT and WDT and the staircase procedure for two-point discrimination thresholds (2PDT), we checked the distributions of the thresholds at the group level. We determined and removed outliers using the same criterion as above (exceeding the range of the third/first quartile +/- 3 IQR). The distributions are shown in Figure S2. There were no outliers in the HPT and 2PDT. In PPT, we removed 2 thresholds (0.7%). In the WDT, we removed 7 thresholds (2.5%).

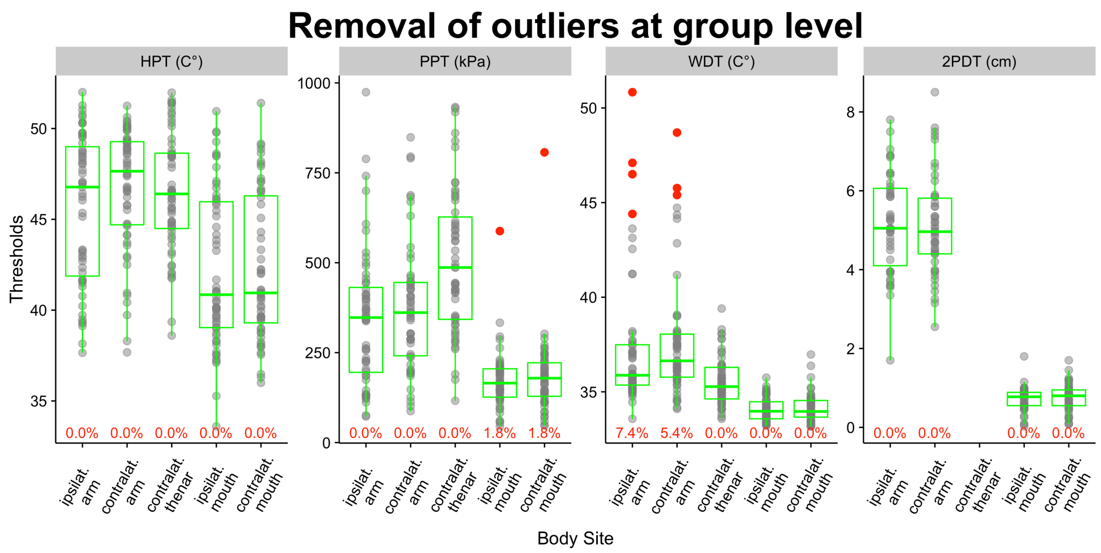


Figure S3: Distributions of heat pain thresholds (HPT), pressure pain thresholds (PPT), warmth detection thresholds (WDT) and two-point discrimination thresholds (2PDT) for all measured body sites (shown on the abscissa) at the level of the entire sample (all groups collapsed). The distributions are shown by green boxplots. Outliers (values exceeding the range between the first quartile -3 interquartile ranges (IQR) and the third quartile + 3 IQR) are highlighted as red points. The proportions of values classified as outliers per body site are shown in red, below the boxplots.

### Assessment of two-point discrimination thresholds—pilot data

To evaluate the procedure for measuring two-point discrimination using the set of calibrated compasses, we performed a pilot. We assessed the within-session test-retest reliability of the absolute threshold attained using the simple staircase procedure and its correlation to the absolute threshold measured using an alternative, more extensive protocol. The alternative protocol consisted in presenting the 23 compasses (ranging in size from 0.7 and 7.3 cm in 3 mm steps) in a randomized order. Compasses between 2.5 and 5.5 cm (middle sized) were presented 10 times, compasses smaller than 2.5 cm (small sizes) or larger than 5.5 cm (large sizes) were presented 5 times. The absolute threshold was determined by fitting a psychometric function to the data. The sample consisted of 10 healthy subjects (6 females, mean age = 29.4, sd = 10.41). All subjects were right-handed and were tested on the right dorsal forearm at 50% distance between the wrist and the elbow. The correlation between two consecutive measurements (T1 and T2) using the staircase procedure was very high (r = 0.81) indicating that the procedure was reliable (see Figure S3 A). The correlation between the thresholds attained using the simple staircase method and the threshold attained using the extensive protocol using randomized presentation and fitting of psychometric functions was also high (see Figure S3 B) which demonstrated convergent validity of the procedures. We concluded that the staircase protocol is an adequate and economic method to measure two-point discrimination thresholds.

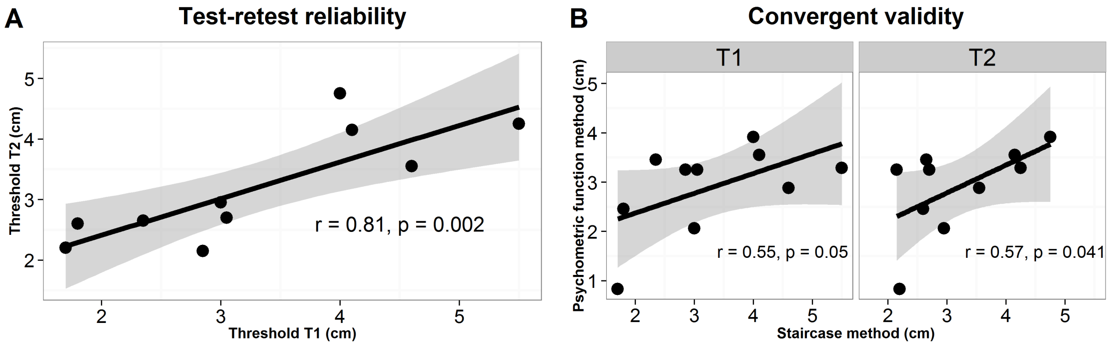


Figure S4: (A) Correlations between two consecutive measurements of two-point discrimination thresholds in 10 healthy subjects using the simple staircase procedure (test-retest reliability). (B) Correlations between the thresholds attained by using the simple staircase procedures with thresholds determined by using a more extensive protocol using randomized presentation of compasses and fitting of psychometric functions (convergent validity). The gray-shaded areas around the regression lines indicate borders of the 95% confidence interval for the linear regression. Pearson correlation coefficients and p-values for the test of a positive correlation are shown in the plots.
